# Not All Predictors Of RCC Clusters Are The Same: An individual sample predictive model to classify patients with metastatic renal cell carcinoma

**DOI:** 10.64898/2026.06.24.734285

**Authors:** Anupama Reddy, W. Jared Brewer, Scott Haake, Brian Rini

## Abstract

Clear cell renal cell carcinoma (RCC) patients have multiple approved therapies, including anti-angiogenesis tyrosine kinase inhibitors (TKIs) and immuno-oncology therapies (IO), but lack a clinically validated biomarker. Using RNAseq data from the IMmotion 151 clinical trial (IM151)^1–3^, 7 RCC biologic clusters have been defined^3^. Several groups have attempted to predict these clusters on various RCC datasets^4,5^ with negative results, suggesting that the association with therapy response from IM151 could not be reproduced. We hypothesized that the specific approaches used to generate cluster predictions led to the misinterpretation of findings. Both published models used standardization (z-scores) to normalize the data within their patient cohorts, imposing an expected gene expression distribution in which ∼50% of patients have apparently higher-than-average expression, artificially impacting the proportion of cluster assignments and leading to potential misclassification. We developed a machine learning (ML) model, IRIS-RCC (Individual RNA-seq Intrinsic Subtyping for RCC), to predict treatment (TKI vs. IO) for patients using an individual-sample model using the IM151 trial (N=823) and validated the model on the JAVELIN Renal 101 trial (JR101; N=726)^3,6^. Our method normalizes gene expression within a given sample using ratiometric expression. This method results in different cluster assignments for individual tumors and distinct clinical correlations. An additional advantage of individual-sample predictions is that they can be readily applied in a prospective setting, where patients must be classified one at a time. IRIS-RCC is currently being validated in a prospective biomarker-driven Phase II clinical trial (OPTIC RCC).

## Methods

The IRIS-RCC machine learning model was developed using RNA sequencing (RNAseq) data from the IM151 trial as the training data (Figure 1A). The original IM151 clusters were consolidated into three functional groups based on treatment associations: (1) Cluster 1+2 (Cluster Angio associated with better response to TKI), (2) Cluster 4+5 (Cluster Immune associated with better response to IO), and (3) Cluster 3+6 (Cluster Exclude with no clear response associations to TKI or IO). Model development utilized a refined subset of 188 genes, consisting of the 25 genes most strongly associated with each cluster, supplemented with manually curated markers.

**Figure 1:**
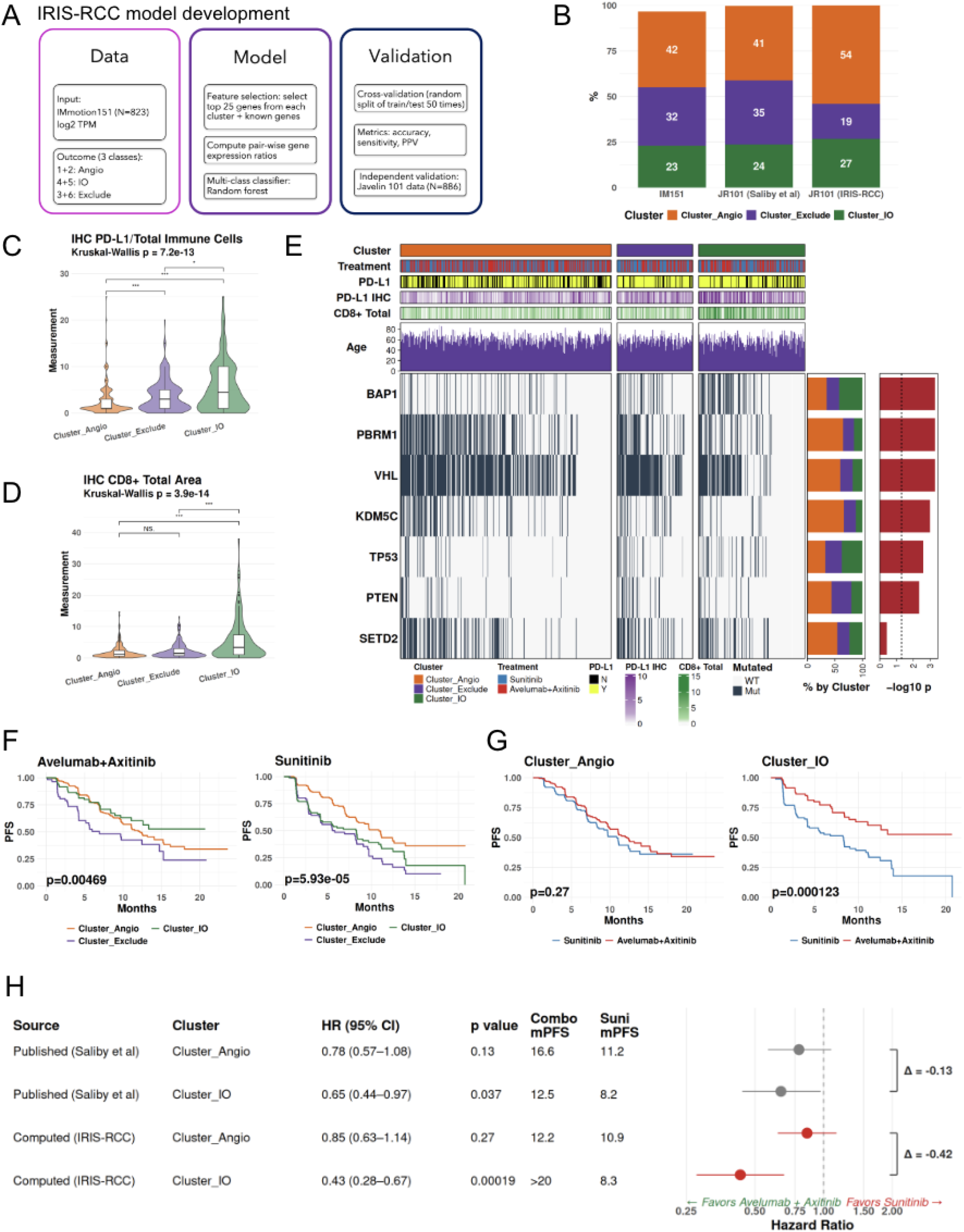
**A**. Conceptual figure showing the modeling used to derive the IRIS-RCC predictor. **B**. Prevalence of the different clusters in the training data (IM151), and validation sets (JR101) using two different methods (Saliby and IRIS). **C**. Violin and boxplots showing the enrichment of PD-L1+ IHC levels as a fraction of the total immune cells in Cluster IO compared to Cluster Exclude and Cluster Angio **D**. Violin and boxplots showing the enrichment of CD8+ IHC levels in Cluster_IO compared to Cluster Exclude and Cluster Angio **E**. Heatmap showing mutational associations with clusters **F**. Forest plot showing hazard ratios for Cluster Angio and Cluster IO for the different treatments across the Saliby and the IRIS-RCC models, statistics by Cox proportional hazards with a Wald test **G**. Kaplan-Meier survival plots for the 3 clusters split by treatment arms, statistics by log-rank p value **H**. Kaplan-Meier survival plots for the treatment split by clusters, statistics by log-rank p value

Rather than using absolute expression, the model employs gene expression ratios to establish relative profiles for each patient. A random forest classifier was then implemented to train the multi-class model. To ensure robust performance, hyper-parameters were optimized through nested cross-validation. The final model was evaluated via bootstrapped cross-validation, repeated 50 times, with performance quantified through accuracy, sensitivity, and positive predictive value (PPV). The resulting cross-validation metrics for the model are 87% for overall accuracy, with sensitivity and PPV of >80% for all three clusters. Cluster Angio, which represented ∼⅔ of the dataset, resulted in the highest cross-validation metrics (95% sensitivity and 91% precision).

## Results

The IRIS-RCC model was validated using the JR101 dataset. Figure 1B shows the frequency of the predicted cluster. We observed a higher proportion of Cluster Angio predictions compared to the other clusters. While comparing the frequencies to the IM151 data and the Saliby et al. predictions with JR101, we observed differences that could be attributed to methodological distinctions, primarily in how gene expression values are normalized^4^. In both previous models, z-scoring was used to perform within-cohort normalization of gene expression values prior to input into the model. This, however, imposes an expectation on the distribution of the expression of any given gene; half of the samples will be given a value of greater than 0, while the other half will be less than 0; this makes each prediction dependent on the patient population in which it was evaluated. Larger, smaller, or differently composed cohorts may yield different predictions for any given sample. IRIS-RCC, by contrast, leverages ratios of gene expression values within a given patient as input features for the model, enabling individual samples to be evaluated in real time, regardless of the overall patient population size or composition. Not only does this facilitate clinical deployment in diverse settings, but it also enables accurate classification of both homogeneous and heterogeneous patient cohorts of any size or composition.

To validate these predictions, we compared statistical patterns observed in clusters from the IM151 data. To reduce circularity in the comparisons, we did not consider comparisons to other features derived from transcriptomics data (e.g., pathway associations), as these are inherently correlated with the gene expression-based clusters. Instead, we focused on associations of the clusters with IHC and mutation data:

1. **PD-L1 IHC association with clusters**. The IM151 study showed that PD-L1+ patients were enriched in Cluster IO and Exclude, but not in Cluster Angio^3^. We observed the same pattern with statistical significance in the validation set (Kruskal-Wallis p < 0.001; Figure 1C). This association was also observed in PD-L1-positive samples (p < 0.001). Note that this association was not reported in Saliby et al.
2. **CD8+ IHC association with clusters**. Cluster IO patients had significantly higher CD8+ IHC levels than Cluster Angio and Cluster Exclude (Kruskal-Wallis test, p < 0.001; Figure 1D).
3. **Mutation associations with clusters**. Consistent with Motzer et al., we found that mutations in key cancer-associated genes were preserved in the JR101 patients based on their cluster assignments (Figure 1E). For instance, *PBRM1* and *KDM5C* mutations are enriched in Cluster Angio, while *TP53* and *BAP1* mutations are enriched in Cluster IO^3^.

Next, we tested our central hypotheses from the training dataset that the predictions from the IRIS-RCC model predict clinical outcome. We expected Cluster Angio patients to have better outcomes with TKIs, while Cluster IO patients would have better outcomes with IO. Indeed, we found that Cluster IO patients had improved PFS with avelumab + axitinib, whereas Cluster Angio patients had better PFS with sunitinib treatment (Figure 1F; log-rank test p = 0.004; log-rank test p = 0.00006). Figure 1G shows a comparison of PFS for patients from a single cluster. We observed that Cluster Angio patients had a similar PFS when treated with either of the TKI-containing therapies, while Cluster IO patients had better PFS with TKI+IO therapy (p = 0.0001; Figure G).

We compared these associations with those previously published by Saliby et al. (Figure 1H), which also trained a random forest model on the IM151 data and validated it on the JR101 cohort^4^. We noted that while the trends were similar, the strength of the association was different, resulting in a larger difference in hazard ratios between Cluster Angio and Cluster IO in our predictions, highlighting the clinical benefit of utilizing the IRIS-RCC as a predictive biomarker. Cluster IO in Saliby et al. was modestly associated with better PFS in the avelumab + axitinib arm (p = 0.03), while we observed a striking difference (p = 0.00001)^4^. Note that the prevalence of the predicted IO group in both Saliby et al. and our model was similar (∼25%)^4^. This highlights the discordance in patient-level predictions between the two methods, which are unable to be precisely assessed with the data available in their publication. The main differences in the two methods are in the processing of the transcriptomics data: Saliby et al. uses z-scores to normalize the data, which is a usual practice for gene expression analysis^4^. However, for ML modeling, this assumes a distribution in gene expression and, therefore, cluster proportions. By utilizing an approach that normalizes expression within each patient, we avoid such assumptions and, instead, enable individual sample-level cluster assignments independent of the trial size or composition.

## Discussion

In this study, we applied IRIS-RCC, a predictive machine learning model that supports the prospective phase II OPTIC RCC, to the JAVELIN Renal 101 study data. As a result of using IRIS-RCC rather than previously published methods that relied on z-scoring normalization for the gene expression data, several key findings emerged. First, IRIS-RCC resulted in individual tumors being reclassified. This is evident when looking at the proportions of tumors classified as Cluster Angio, IO, or Exclude in the JAVELIN Renal 101 dataset. Using the IRIS-RCC classifier, relatively more tumors were classified as Cluster Angio and IO and fewer tumors were classified as Cluster Exclude. As a result of this reclassification, we observe a more robust benefit of IO therapy in the Cluster IO patients. This study functions as an important external validation of the gene expression signatures identified originally in the IMmotion 151 study^3^.

A key finding in this study is that choice of methods are an important consideration when applying gene expression signatures (i.e., clusters) retrospectively to datasets. The same tumor can be assigned to different gene expression clusters depending on the methods, such as z-scoring versus gene ratios for normalization of gene expression data. As a result, correlations between treatment outcomes and gene expression-defined clusters can differ in a methods-dependent manner. Not all references to the IMmotion 151 clusters are using the same methodology and thus not all clusters are the same. Methods that assume that the relative proportions of gene expression-defined tumor groups are nearly identical between distinct cohorts should not be used. Just as two cohorts may contain different proportions of clinically-defined prognostic subgroups (e.g., within the context of clear cell renal cell carcinoma, trial A may enroll more IMDC favorable risk patients than trial B), it is also plausible that two distinct trials may contain different proportions of gene expression-defined tumor groups. One advantage of the gene ratio method for gene expression normalization utilized in the current manuscript is that it does not assume these proportions are the same across clinical trial cohorts.

This current analysis has several limitations. It is a retrospective analysis and does not obviate the need for prospective validation of the IMmotion 151 signatures. Rather, it supports the prospective OPTIC RCC trial by validating the IMmotion 151 signatures retrospectively in an additional cohort. Ideally, a prospective study that randomizes patients with the same tumor gene expression profile to different therapies is needed to prove which therapy is best for an individual tumor type. Furthermore, the majority of tumor gene expression data from the JAVELIN Renal 101 study was derived from primary tumors. Gene expression signatures in primary tumor and metastatic sites are often discordant. Indeed we observed 40% discordance in samples obtained as part of the OPTIC trial^7^.

## Summary

In summary, this study demonstrates that patients with metastatic RCC enrolled in Javelin Renal 101 and have the Cluster IO gene expression tumor signature have superior outcomes when treated with the IO-containing regimen. Meanwhile, Cluster Angio patients have similar PFS when treated with either the TKI or IO/TKI arm, since both contain an anti-angiogenic therapy. Z-scoring should not be used to predict clusters as it leads to erroneous conclusions by account of its context-dependent nature and variation-over-time effects. Focusing methodology on those approaches able to deterministically predict samples independently from one another will enable robust, reproducible, and clinically-relevant patient stratification and therapeutic assignment.

